# Interindividual differences (bold and shy strategies) in neophobia may explain the invasive success of the worm *Eisenia fetida*

**DOI:** 10.1101/2025.01.10.632465

**Authors:** Alexandra Rodriguez Pedraza

## Abstract

When they are confronted to novel environments individuals may express different kinds of behaviors. They can move a little or even stay immobile with few exploration of the new environment or they can increase their activity, move a lot and explore actively the new place. Several studies on neophobia have been conducted on vertebrate species and they distinguish bold and shy patterns in animals’ reactions. Less studies have been conducted on invertebrates in this area. Here I present the case of the worm Eisenia fetida that I tested in open field tests in order to detect if they present different response profiles when confronted to a novel environment and how these profiles can vary depending on size factor. I was able to distinguish two profiles, a shy/philopatric reaction present in young and adults and a bold/explorer reaction that can be observed in mature worms. From the 311 worms tested one half was bold/explorer and the other half was shy/philopatric. The existence of these two profiles may explain the invasiveness of the species: some indivivuals stay and occupy the known environments and some individuals enhance their activity in order to colonize other environments.

## Introduction

Psychologists and ethologists (Bates 1986, Chapple et al. 2012) have defined temperament and personality, two concepts that are more and more used to describe the interindividual differences in the expression of several behaviors present early in individuals’ lives (Bates 1989, Goldsmith et al. 1987). Temperament depends on the genetic characteristics of the individuals and personality is the result of environmental effects on temperament. Animal personality can be defined as consistent differences in the behaviour of individuals across time and situations (Sih et al. 2004) Personality and temperament can include aspects of behavioural responses as reactions towards predators, neophobia and social behaviour towards conspecifics. These behavioral characteristics may condition dispersal behavior, approach towards novel objects and novel food items consumption (Sinn et al. 2008).

The literature concerning interindividual differences in personality is very extensive and some studies converge to say that constancy exists in species like horses and salamanders (Le Scolan et al. 1997, Sih et al. 2003) whereas other studies found that there is no constancy across the contexts in species like pumpkinseed sunfishes and prairie voles (Coleman and Wilson 1998, Neff and Sherman 2004, Lee and Tang-Martinez 2009). These different observations vary depending on the studied species, on the methods that are used and on the behavioural aspects that are explored. Even if invertebrate species represent over 95% of all animal species (Scheffers et al. 2012, Brusca and Brusca 1990), personality studies have traditionally focused on vertebrate species (Gosling 2001). As they present higher cognitive abilities, vertebrates were considered as more likely to exhibit complex behaviours while invertebrates were thought to behave stereotypically, thus showing few or no individual differences in their behaviour. Recently, more efforts have been made to include invertebrates in personality research (Kralj-Fišer and Schuett 2014; Mather and Logue 2013). These studies were mainly focused on a few taxa in molluscs and arthropods. Cephalopods which are considered as advanced invertebrates considering cognitive functions have been more studied in terms of personality than most groups of invertebrates (Sinn et al. 2005, Carere 2015). However, it is being increasingly recognised that the study of personality in invertebrates is of particular interest, as compared to that of vertebrates, given their ecology and biology. In addition, they are easier to rear and to maintain in laboratory conditions. Finally, their short life cycles make them good study models.

Neophobia is one of the aspects of personality and it consists in the fear towards novel stimuli or novel situations. A stimulus can be novel because it has simply never been experienced by an individual during its lifetime (Brown et Chivers 2005). Neophobia has recently received much ecological attention, primarily in the context of decision making and even in large scale behaviors as migration. For example, low levels of neophobia were reported in migratory Sardinian warblers, *Sylvia melanocephala momus*, when compared to garden warblers, *Sylvia borin* (a non-migratory species) which behave as philopatric individuals (Mettke-Hofmann et al. 2005). Furthermore, in a study conducted by Candler and Bernal (2015), differences of boldness were observed in cane toads. Actually, in this species individuals from native populations did not approach a novel object while more than half of the individuals from introduced populations did. It has also been reported that early experience with novel objects in laboratory environments can result in low neophobia levels in young hand reared parrots *Amazona amazonica* compared with individuals raised by their parents in the simpler nest box environments that present lower objects diversity (Fox and Millam 2004). Neophobia towards a specific situation can be measured only one time as the situation can be considered as novel only the first time the animal is tested in.

As we saw above, personality and temperament have been studied in several vertebrate species and it has been observed that different factors influence their expression, from genes to environmental conditions (Tremmel and Muller 2013). Personality has been studied in invertebrates as the crayfish *Pacifactacus leniusculus* (Galib et al. 2022), and in beetles (Labaude et al. 2018) while neophobia has been studied in ants (Rodriguez 2023) but none of these aspects has been tested in worms. In ants it has been demonstrated that there are far explorers and close explorers that differ in the distances they move from the nest and it has been observed that far explorers are more mobile than close explorers. In many species two types of individuals have been described concerning migration or movement status. Actually, it has been demonstrated that there are philopatric and dispersing individuals that can be distinguished because dispersing individuals move to new environments to reproduce while philopatric individuals stay in the original region and reproduce in their original known habitat. The study of personality in invertebrates could provide a better understanding in other fields, such as population and community ecology (Modlmeier et al. 2015, Wolf and Weissing 2012). Recent studies suggest that aspects of personality as neophobia could affect species distributions, species invasions and response to environmental changes (Sih 2004, Chapple et al. 2012).

Worms are one of the most represented taxa in the world. This group contains a huge number of species and it is present in several soils across the world (Brown et al. 2013). The success of worms in the world is mainly due to their social organisation and their reproductive system. Actually, worms live in big groups and reproduce by a hermaphrodite and parthenogenetic ways that facilitate the renovation of generations (Diaz Cosin et al. 2010). This aspect is reinforced by the fact that they are animals with a very short life cycle. However, no study has focused on their behaviour and no study has explored the personality or the neophobia in worms. In this study we hypothesized that worms present a high diversity of reactions to novel contexts that confer them the possibility to colonize both familiar and unfamiliar environments.

## Material and methods

### Experimental protocol

Open field tests have been used from decades to study exploration in vertebrates like rats. These tests measure exploratory locomotor behaviour and general activity in rodents. Since its introduction, 40 years ago, the open-field test has become one of the most widely used instruments in animal psychology to test aspects of personality. The open field is an enclosure generally square, rectangular or circular (Gould 2009).

Originally introduced as a measure of emotional behaviour in rats, open field test has proven to be equally successful with mice. The test offers the possibility to assess novel environment exploration, general locomotor activity, and some aspects of behaviour in rodents. When placed in an open field, animals may display different behaviours. Some wander through the area to explore the new environment. Rodents will typically spend a significantly greater amount of time exploring the periphery of the arena, usually in contact with the walls than the unprotected centre area (Taiwo 2015). Open spaces induce fear behaviour in animals that have an aversion to unfamiliar environments, and these animals will seek dark regions or protective crevices. Rodents that spend significantly more time exploring the unprotected centre area are considered bolder and less anxious than shy individuals that explore in close contact with the walls.

In this study I adapted the classical open field protocols from rodents studies to the study of woms’ neophobia towards a new environment.

For this aim I tested 311 worms from *Eisenia fetida* species.

### Biological material

Worms are animals that belong to 13 families, and, even if more than 5000 have been described (Brown et al. 2013), *Eisenia fetida* is the most used for vermiculture because this species is able to convert almost any kind of organic waste into a final product called vermicompost (Decaens et al. 2003). These organic fertilizers improve the soil chemical proprieties as well as it physical and biological proprieties (Martínez 2009). Worms conduct an important activity in the environment (Oliveira et al. 2008). This activity consists in the production of humus (Durán and Henríquez 2009) because its excrements are an excellent organic fertilizer with the content of its bacterial flora (Toccalino et al. 2004) and they have the ability to transform organic material into nutrients (Ávila et al. 2007) that combine with soil and improve its productivity (Mendoza 2018).

*Eisenia fetida* is a worm species very common and very used for the recycling of organic waste thanks to the vermicompost. It is also very used for ecotoxiological, physiological and genetic studies. This extended use is due to its ubiquitous and cosmopolitan distribution with short life cycles, a big range of tolerance to temperature and humidity and an easy and simple management (Domínguez 2004). *Eisenia fetida* corresponds to the stripted morphotype which has a non-pigmented area between the segments **(Figure 1)** or a pale yellow area (Dominguez and Perez Losada 2010). They live in mixed colonies in manure piles and vegetal waste.

**Figure 1.**
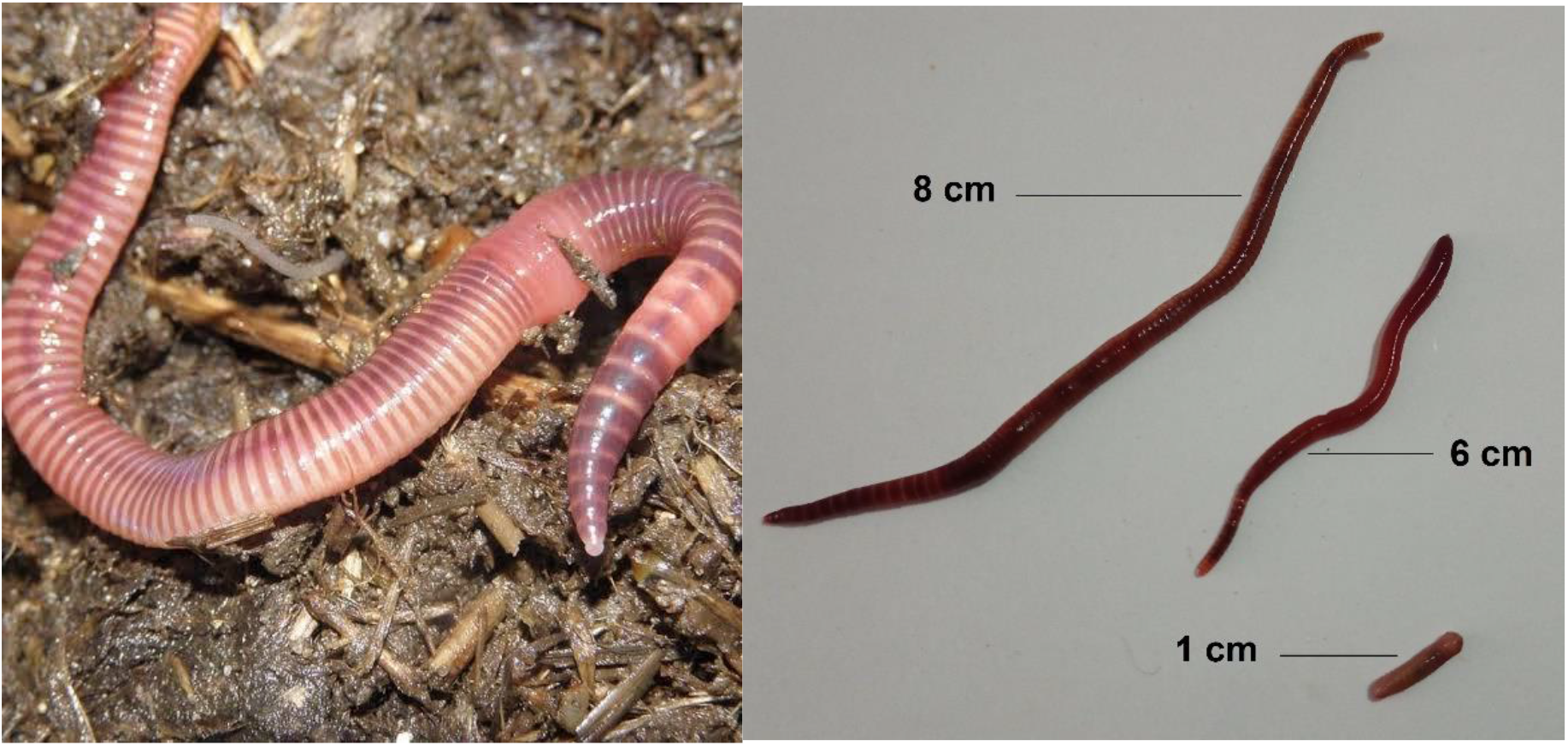
a. *Eisenia fetida* b. *Eisenia fetida* of different sizes.

311 *Eisenia fetida* worms were reared in a urban vermicompost in Bogotá (Colombia) in order to conduct the experiments described bellow.

### Experimental procedure

For each test I put the worm on a clean plastic table and I recorded its behaviour during 2 minutes. Each worm was video recorded during the whole test.

The experimental area was a rectangular field of 27,5cm x 21 cm divided into 15 areas as bellow in **Figure 2**.

**Figure 2.**
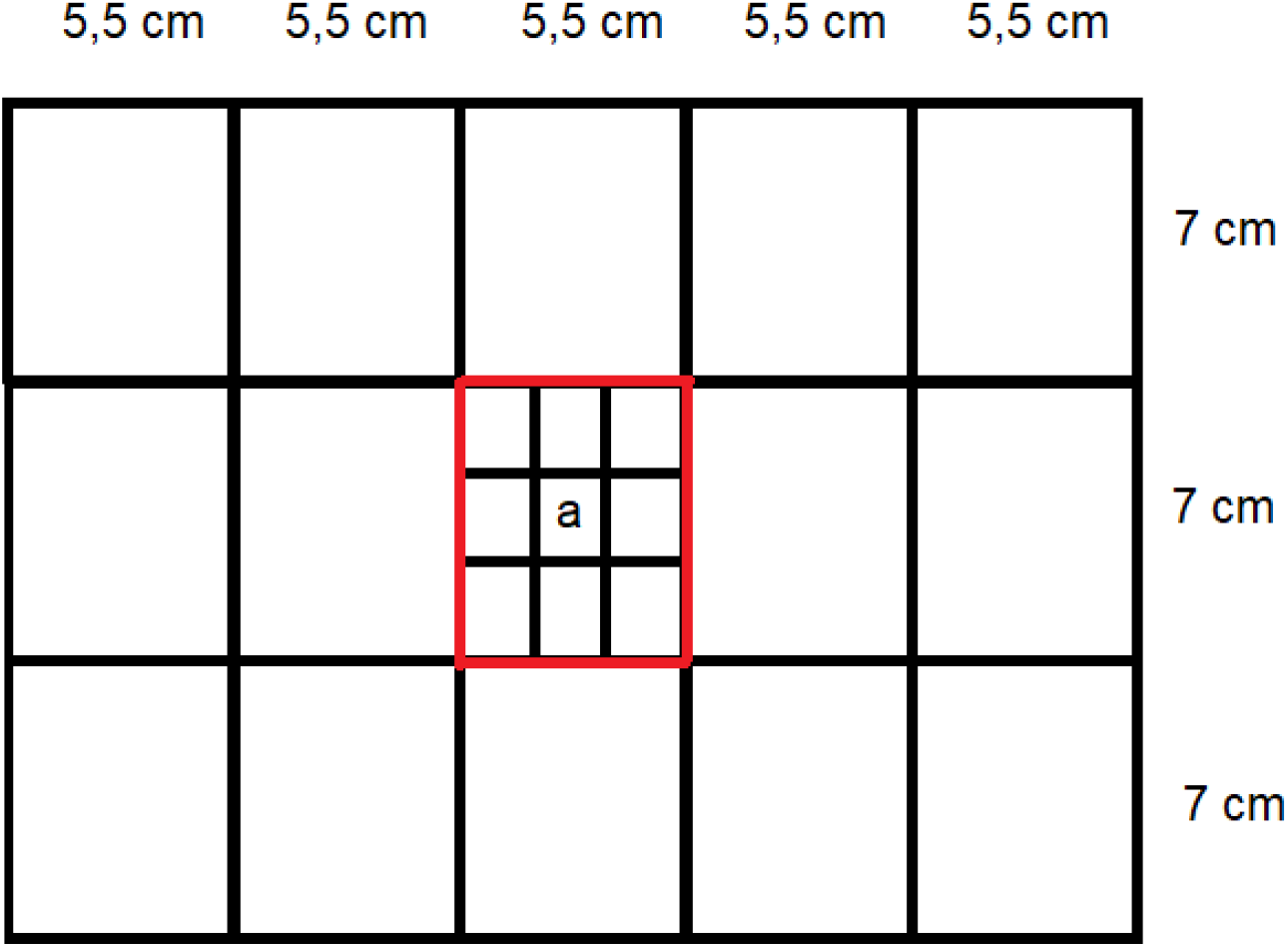
Experimental apparatus: rectangular plastic table.

For the experiment each worm was put into the little central cell a. Time to get out from the central red cell was recorded. When a worm did not leave the central cell during the whole experiment, I scored the value 200 for the time.

The number of visited big cells was noted as well as the number of little cells of the centre of the experimental area. If the worm visited one big cell and one little cell, I noted that 1,1 cell was visited. If the worm visited 3 big cells and 2 little cells, I noted that 3,2 cells were visited.

Worms make elongations of their body followed by a body contraction and another elongation and then successively in order to advance. I noted the number of elongations conducted during the whole experiment.

All the experiments were conducted in Bogotá from 4/12/2024 to 6/12/2024 at a stable temperature comprised between 19 and 20°C.

### Statistical Analysis

I conducted linear regressions between the number of visited cells and the worms’ size and between the number of elongations and the worms’ size.

Two types of worms were identified; the ones that never left the central red cell that were considered as shy/philopatric worms because they stayed in the original area and the ones that visited at least one big cell different from the original one. These last worms were considered as bold/explorers as they left the original cell to explore other cells.

The worms were divided into 8 classes depending on their size as in **figure 5**. I conducted Mann-Withney tests in order to compare the number of elongations between the philopatric and the explorer worms of each size class to know if bold/explorer worms were more mobile than shy/philopatric ones.

I conducted a linear regression between the number of visited cells and the time to exit the central cell in order to know if the individuals that visited more cells were also the ones who left earlier the central area.

## Results

The number of visited cells was positively and significantly correlated to the worms size (R^2^ = 0,19 p< 0,001) indicating that the larger the worms were the more cells they visited **(Figure 3)**.

**Figure 3.**
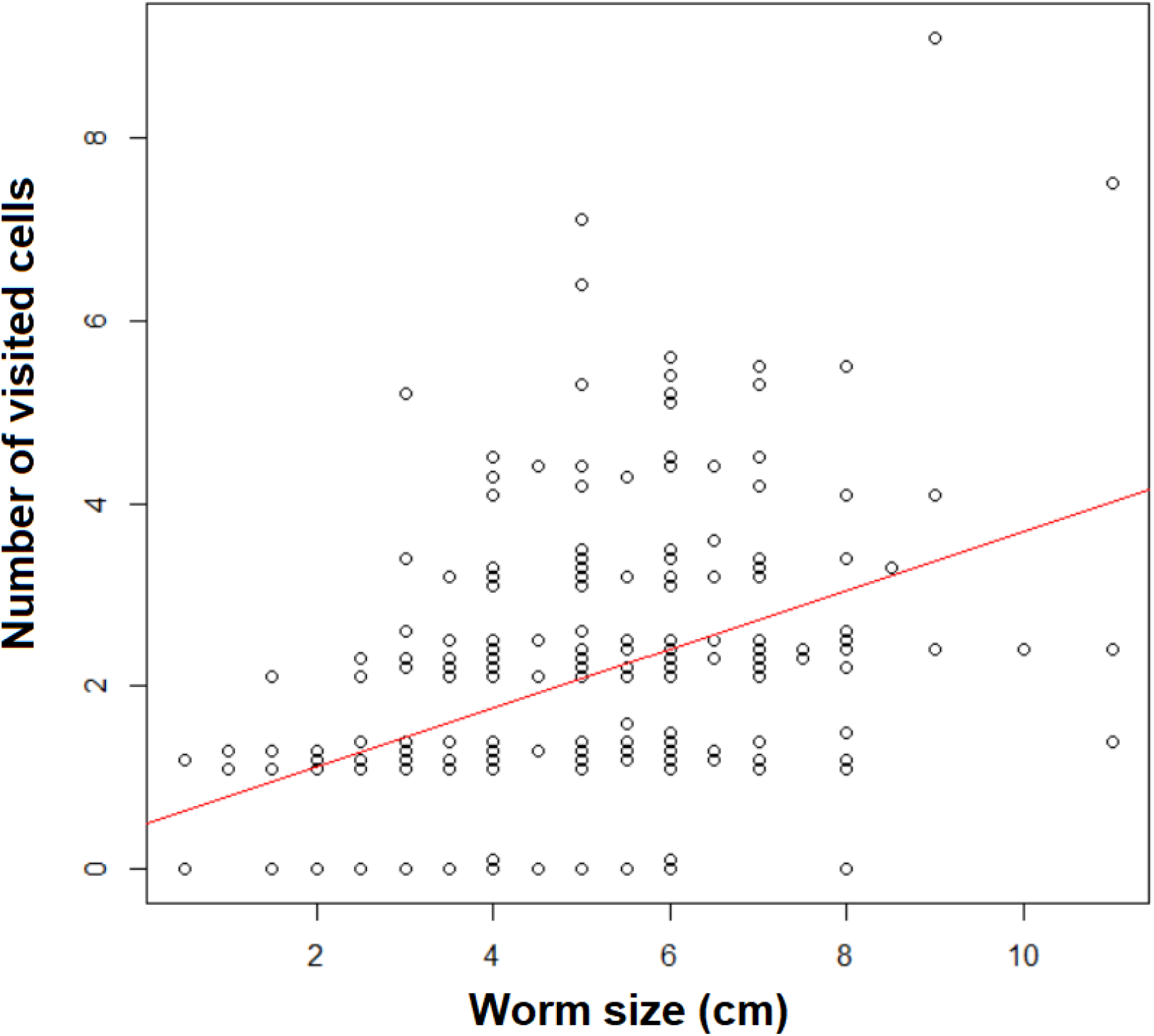
Number of visited cells versus worm’s size.

The number of elongations was positively and significantly correlated to the worm size (R^2^=0,02, p <0,01) indicating that the larger the worms were the more they were mobile **(Figure 4)**.

**Figure 4.**
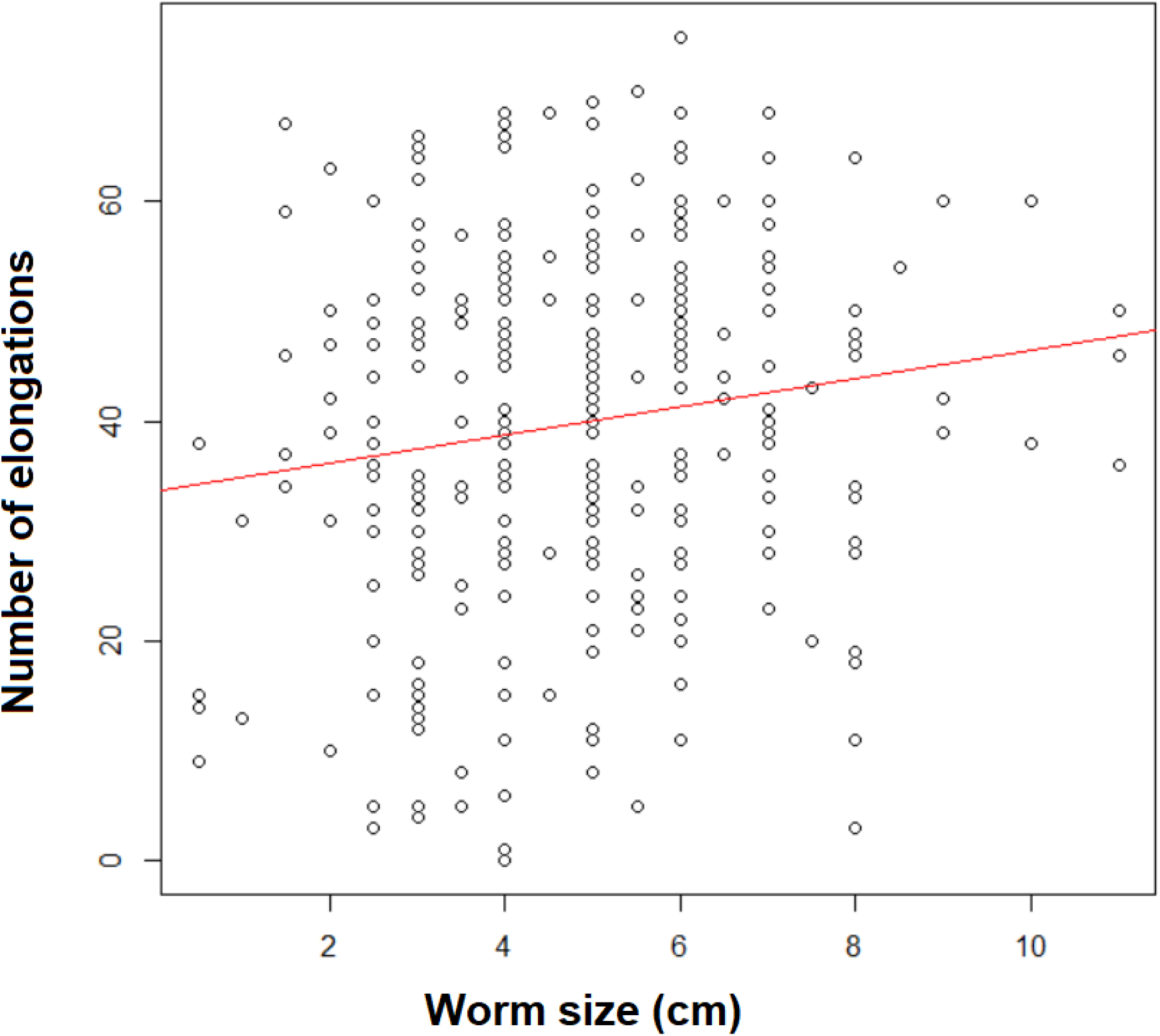
Number of elongations versus worm’s size.

147 worms displayed shy behaviour and did not leave the central cell. 164 worms displayed bold behaviour and left the central cell.

For each size class except for the two first classes (from 0 to 2,5 cm) there were bold/explorer and shy/philopatric individuals. For the two first classes there were only shy/philopatric individuals **(Figure 5)**

**Figure 5.**
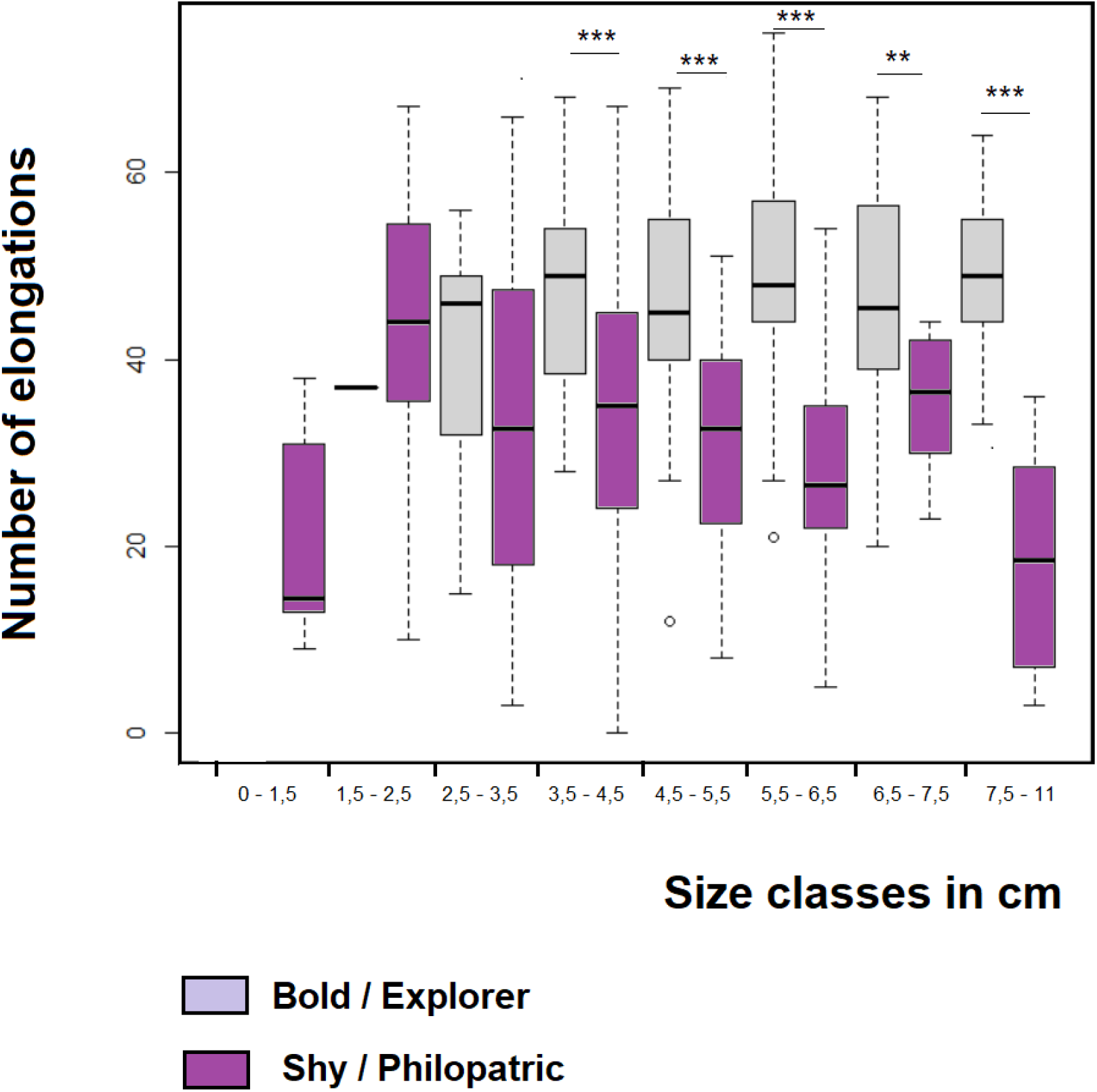
Number of elongations displayed by the worms of each size class.

For the size class including worms of 2,5-3,5cm there was no difference in the number of elongations between shy/philopatric and bold /explorer worms. For the other classes explorer individuals were significantly mor mobile than philopatric ones.

Mann Whitney test p values: * : p<0.05 * *: p<0.01 ** *: p<0.001

The number of visited cells was negatively and significantly correlated to the time to exit the central cell (R^2^=0,59 p<0,001) indicating that the more the worms left early the central cell the more the number of visited cells was high **(Figure 6)**. That means the high explorer worms were also early explorer worms.

**Figure 6.**
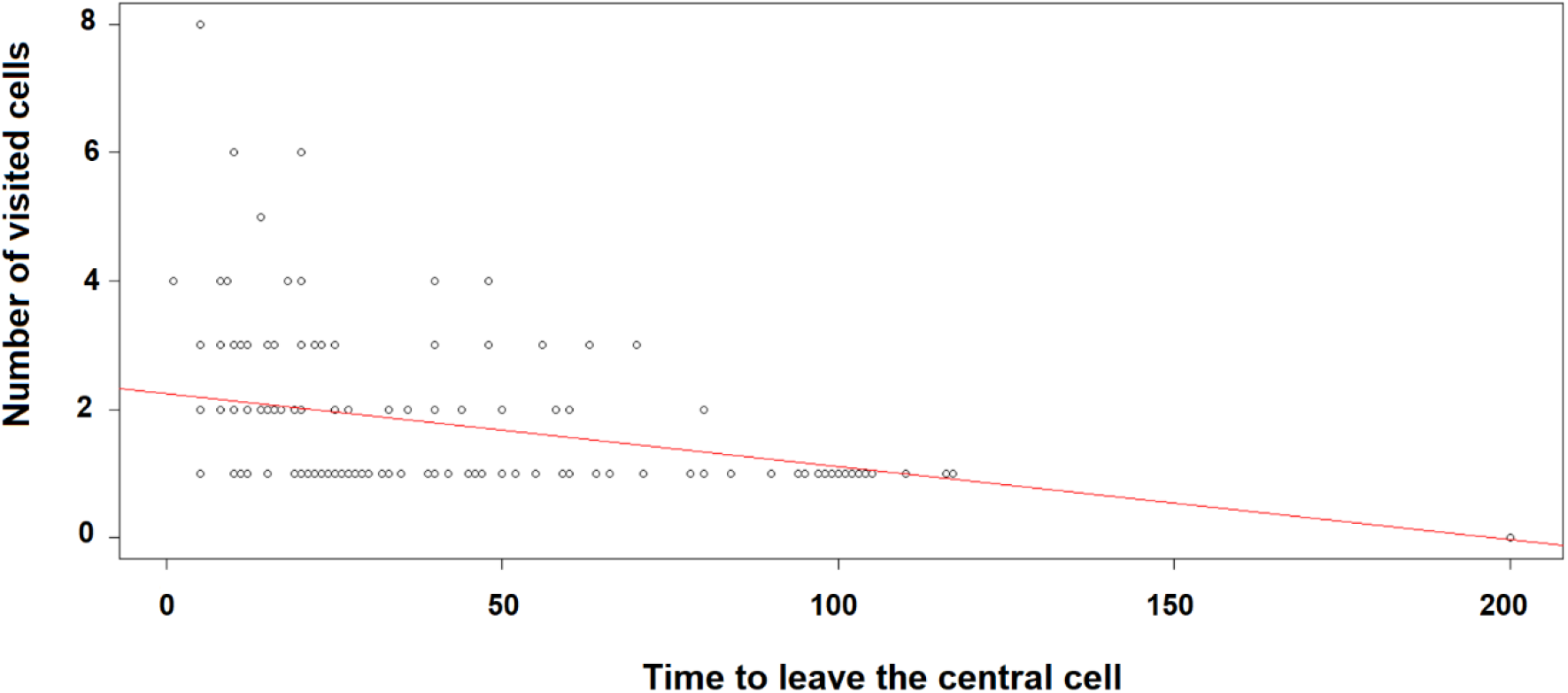
Number of visited cells versus time to leave the central cell.

## Discussion

In this study I found for the first time that differences in neophobia in an open field test can be observed between worms from a same social group of three hundred worms. There were worms very mobile and worms who displayed few movements during the experiments. There was also a big diversity in time to exit the central cell as some worms left the cell one second after the beginning of the experiment and other worms did not leave the central cell during the whole experiment.

I observed that the number of visited cells was positively correlated with the worms’ size which seems to indicate that levels of exploration are directly related to the size, thus to the maturity and age status of the worms. Actually, worms size depends on their age (Schuldt et al. 2005).

The number of elongations was also positively correlated to worm size which indicates that older and larger worms move more than younger ones. This result is different from the one observed in birds as it has been demonstrated that young starlings move more than adults (Rodriguez et al. 2021a). By consequence we can consider that dispersal strategies depend on the considered species: in some cases, dispersal will be ensured by young individuals and in other species it will be conducted by adult or old individuals.

Concerning the type of profiles detected, I observed that half of the worms were explorer and the other half were philopatric. There were explorer and philopatric worms in all the size classes except for the two first classes. That means that in worms there are individuals that spread easier than others probably because they display different reactions to novelty: bold and shy reaction. The existence of shy/philopatric individuals and bold/explorer individuals may explain why worms have colonized several soil habitats in the world. In particular it can explain why *Eisenia fetida* has become an invasive species as there are individuals who stay in their original habitats and individuals that move towards new ones.

These two types of profiles are well defined when worms are longer than 3,5 cm. Between 2,5 and 3,5cm there are no significant differences in the number of elongations between philopatric and explorer worms. This is probably due to the fact that they are not mature enough: clear patterns of behaviour are probably well shaped after this stage.

In a study conducted by Schuldt and collaborators (2005) it has been demonstrated that *Eisenia fetida* reaches sexual maturity when it reaches 0,25g which corresponds to 2,5-3 cm. We found that the size classes under 2,5cm did not contain explorer individuals but only philopatric ones. This last observation indicates that high mobility is reached at sexual maturity in worms, by consequence they start to disperse and explore more when they are able to reproduce. Under this stage individuals are too young or little and belong to a prior shy phase and individuals who reach sexual maturity are big enough to disperse or they may be hormonally motivated to disperse and reproduce. We can think that explorer individuals over 2,5 cm are the ones responsible of reproduction and dispersion in the soil. They probably move to other places in order to colonize new areas or to reproduce with other individuals by the hermaphrodite way and exchange genetic material contained in sperm. However, more studies should be conducted as I tested the worms on a plastic tale. Tests in earth conditions could bring more information on worms’ dispersion in the soil. Our observations can apply to the moments when worms reach the surface of the soil and disperse over this area.

The more the worms were large the more they displayed high levels of mobility. This observation means that new habitats are probably colonized by the larger worms which may lead colonization processes. This observation is in accordance with the observations in ants (Rodriguez 2023). Actually, soldier ants which are bigger than workers explore and move more in an open field experiment.

In this study I also observed that the number of visited cells was negatively correlated to the time to exit the central cell. By consequence we can deduce that bold individuals are not only individuals who visit more cells but they are also early explorers.

A relation between neophobia and dispersal has been observed in vertebrates like birds (Great tits, *Parus major*) (Dingemanse et al.2003) and in terrestrial tortoises like Hermann tortoise, *Testudo hermanni*. (Rodriguez et al. 2021b) Individuals that are bold in neophobia tests travel higher distances than shy ones in the wild (Dingemanse et al.2003, Rodriguez et al. 2021b). Moreover, Dingemanse and collaborators (2003) observed that fast exploring great tits had offspring that dispersed further. They also obtained that immigrant individuals arriving in a new habitat, were faster explorers in novel environment tests than locally born individuals. Here I observed for the first time that an invasive worm species comprises individuals with different tendencies to disperse; those who explore and disperse further are also the ones who move the more and who leave earlier the central cell in the open field test.

In ants, the results of the comparison between five species also suggest that they differ in their reactions towards a novel environment. In this taxon, the species that tend to colonize more ecosystems and that are settled in open spaces like urban areas, islands and rocky landscapes more exposed to the sun and to predators appeared to be bolder and more active in the open field test. Here we observed that there were two kinds of profiles in a same species which live under the surface of the soil. We can think that both in ants and in worms that are underground animals the biggest individuals contribute to dispersal and to the construction of tunnels as they are more explorer and mobile. Large worms can be good candidates for aerating the soil in different directions by tunnel construction and to digest waste in the soil. They can be used for better vermicompost functioning.

Jones and Gray observed behavioural differences between males and females in an open field test that evaluates neophobia. In one hand, Jones (1977, 1982) observed that female chicks presented less behavioural inhibition when placed in a novel environment or in open field than males. In the same way, female rodents seem to explore sooner a novel environment than males (Gray 1971, 1974). In the other species like ungulates, primates and lepidopters,, females appear to be more fearful than males (Buirski et al. 1978, Crepeau and Newman 1991, Rodriguez 2025). Actually, in a lepidopter species like Sangalopsis microleuca males are bolder and move more than females in an open field test. Here I studied a hermaphrodite species and I found that there were both bold/explorer and shy/philopatric individuals in all the adult classes. Thus, there are two different strategies in a species in which individuals act sometimes as males and sometimes as females. This diversity of behaviours combined with the hermaphrodite strategy may explain the success of this species in the world.

In invertebrates and in particular in insects the individuals increase their activity with temperature (Abdullah 1961, Neven 2000, Rodriguez 2023). Here I conducted the experiments in a stable environment where temperature was comprised between 19 and 20°C. More studies should be conducted with variable temperatures in order to know if worms’ mobility varies with this factor.

Different debates have been raised and major questions are still open concerning three essential points. One first question is if there is constancy or repeatability of interindividual differences across the time (Jones 1977). In one hand, constancy across the time of interindividual differences is generally interpreted as the existence of temperaments or personalities (Gosling 2001). On the other hand, the absence of consistency is interpreted as the expression of context or state dependent behaviours (Van Oers et al. 2005) or as the expression of behavioural plasticity (when the individual can behave in one way one time and in another way another time) (Pfennig et al. 1993, Neff and Sherman 2003). More studies on invertebrate interindividual differences must be conducted in order to know if they are really different personality stable traits. In my study I tested the worms only one time as I was interested by neophobia. Successive tests repeating the same protocol could be informative to know if the worms behave in similar ways in all the tests and to confirm if personalities exist in this species.

Greenberg (2003) proposed that the primary benefit of neophobia is protection against danger in the Dangerous Niche Hypothesis (where neophobia increases with the level of danger in the niche). Neophobia can be driven by natural selection based on environmental patterns because neophobia can have a strong innate (genetic) component (Mettke-Hofmann, 2017). However, individual neophobic responses are also shaped by ontogenetic experience, which can result in more or less neophobia (phenotypically plastic neophobia: Brown et al. 2013). More studies should be conducted in order to better understand how this neophobia may has been modulated in space and time during the evolution of the species and how it can explain the amazing adaptations of worms to several ecosystems. Actually, neophilic individuals (who are attracted towards novel stimuli) may be responsible of colonization processes in novel environments and neophobic ones may be the individuals that ensure the maintenance of species in their original habitats.

